# Multifactorial Chromosomal Variants Regulate Polymyxin Resistance in Extensively Drug-Resistant *Klebsiella pneumoniae*

**DOI:** 10.1101/134684

**Authors:** Miranda E Pitt, Alysha G Elliott, Minh Duc Cao, Devika Ganesamoorthy, Ilias Karaiskos, Helen Giamarellou, Cely S Abboud, Mark AT Blaskovich, Matthew A Cooper, Lachlan Coin

**Author notes:** Address correspondence to authors:Lachlan Coin & Matthew A Cooper.

## Abstract

Extensively drug-resistant *Klebsiella pneumoniae* (XDR-KP) infections cause high mortality and are disseminating globally. Identifying the genetic basis underpinning resistance allows for rapid diagnosis and treatment. XDR isolates sourced from Greece and Brazil, including nineteen polymyxin-resistant and five polymyxin-susceptible strains, underwent whole genome sequencing. Approximately 90% of polymyxin resistance was enabled by alterations upstream or within *mgrB*. The most common mutation identified was an insertion at nucleotide position 75 in *mgrB* via an ISK*pn26*-like element in the ST258 lineage and ISK*pn13* in one ST11 isolate. Three strains acquired an IS1 element upstream of *mgrB* and another strain had an ISK*pn25* insertion at 133 bp. Other isolates had truncations (C28STOP, Q30STOP) or a missense mutation (D31E) affecting *mgrB*. Complementation assays revealed all *mgrB* perturbations contributed to resistance. Missense mutations in *phoQ* (T281M, G385C) were also found to facilitate resistance. Several variants in *phoPQ* co-segregating with the ISKpn26-like insertion were identified as potential partial suppressor mutations. Three ST258 samples were found to contain subpopulations with different resistance conferring mutations, including the ISKpn26-like insertion colonising with a novel mutation in *pmrB* (P158R), both confirmed via complementation assays. We also characterized a new multi-drug resistant *Klebsiella quasipneumoniae* strain ST2401 which was susceptible to polymyxins. These findings highlight the broad spectrum of chromosomal modifications which can facilitate and regulate resistance against polymyxins in *K. pneumoniae*.

**DATA SUMMARY:**

1. Whole genome sequencing of the 24 clinical isolates has been deposited under BioProject PRJNA307517 (https://www.ncbi.nlm.nih.gov/bioproject/PRJNA307517).

**IMPACT STATEMENT:** *Klebsiella pneumoniae* contributes to a high abundance of nosocomial infections and the rapid emergence of antimicrobial resistance hinders treatment. Polymyxins are predominantly utilized to treat multidrug-resistant infections, however, resistance to the polymyxins is arising. This increasing prevalence in polymyxin resistance is evident especially in Greece and Brazil. Identifying the genomic variations conferring resistance in clinical isolates from these regions assists with potentially detecting novel alterations and tracing the spread of particular strains. This study commonly found mutations in the gene *mgrB*, the negative regulator of PhoPQ, known to cause resistance in KP. In the remaining isolates, missense mutations in *phoQ* were accountable for resistance. Multiple novel mutations were detected to be segregating with *mgrB* perturbations. This was either due to a mixed heterogeneous sample of two polymyxin-resistant strains, or because of multiple mutations within the same strain. Of interest was the validation of novel mutations in*phoPQ* segregating with a previously known ISK*pn26*-like element in disrupted *mgrB* isolates. Complementation of these *phoPQ* mutations revealed a reduction in minimum inhibitory concentrations and suggests the first evidence of partial suppressor mutations in KP. This research builds upon our current understanding of heteroresistance, lineage specific mutations and regulatory variations relating to polymyxin resistance.

## INTRODUCTION

*Klebsiella pneumoniae* (KP) strains classified as extensively drug-resistant (XDR) are rapidly emerging due to the dissemination of plasmid-encoded resistance towards aminoglycosides, β-lactams, fluoroquinolones and carbapenems [1]. Notably, carbapenem-resistant KP have been linked to high morbidity and an overall mortality of 48% in infected patients[2]. Polymyxin B and colistin (polymyxin E) are now one of the last viable therapeutic options [3]. Unfortunately, resistance to this last-line antibiotic class is an increasing global burden, with countries particularly impacted including Asia (Korea [4, 5], India [6, 7]), Europe (Greece [8–10]), Italy [10, 11]) and Latin America (Brazil [12, 13]). There is considerable debate regarding the mortality associated with polymyxin-resistant infections. Combining several clinical cohorts has provided an overall mortality estimate ranging from 20 to 100% which was dependent on early detection of the outbreak [14].

Polymyxins infiltrate Gram-negative bacteria via initial binding to the basal component of lipopolysaccharide, lipid A. This causes the displacement of Mg^2+^ and Ca^2+^, disrupting bacterial outer membrane integrity, allowing the polymyxins to traverse the inner membrane and act on intracellular targets. An extended exposure in KP triggers the activation of the two-component regulatory systems, PmrAB and PhoPQ [15–17]. These systems regulate a pathway that modulates *pmrC* and the*pmrHFIJKLM* operon facilitating the addition of phosphoethanolamine (pEtN) and/ or 4-amino-4-deoxy-L-arabinose to lipid A phosphate groups, resulting in impaired polymyxin binding interactions [18–20]. Disruption of *mgrB*, the negative regulator of PhoPQ, has been commonly observed in isolates of clinical origin [8, 21]. The constitutive up-regulation of *pmrC* and the *pmrHFIJKLM* operon incurs a minimal fitness cost and appears to be stable, with minimal reports of reversions [22, 23]. Heteroresistant populations, where only a subset of bacteria are resistant, have been reported in KP which complicates diagnosis [24]. The emergence of pandrug-resistant KP is of grave concern [25] and this acquisition of resistance is further exacerbated by the recently reported plasmid-encoded colistin resistance gene *mcr-1*, which encodes a pEtN transferase enzyme, albeit currently rare in KP [26].

This study aimed to investigate XDR-KP clinical isolates arising in Greece and Brazil during 2012 to 2014 to identify and validate genetic variants contributing to resistance. These alterations were compared to prior clinical isolates to ascertain if these mutations have been previously detected globally.

## METHODS

### Bacterial isolates

KP clinical isolates were acquired from the Hygeia General Hospital, Athens, Greece and Instituto Dante Pazzanese de Cardiologia, Brazil from patients in 2012 to 2014. Cultures were supplied as stabs/slants or on agar, and were subsequently cultured in Nutrient Broth. Cultures were made to 20% (v/v) glycerol and stored at −80 °C. When required for assay or extraction, glycerol stocks were struck out to obtain single colonies on either Nutrient Agar or Tryptic Soy Agar with 5% defibrinated sheep blood. Reference strains included *Escherichia coli* (ATCC 25922) and *Klebsiella spp*. (ATCC 13883, ATCC 700603, ATCC BAA-2146), which were obtained from the American Type Culture Collection (ATCC; Manassas, VA, USA).

### Antimicrobial susceptibility assays

Species identification and susceptibility profiles of clinical isolates from Greece and Brazil were evaluated in the clinic using VITEK^®^2 (bioMérieux). Strains were further validated at the Institute for Molecular Bioscience (IMB) (The University of Queensland, Australia) using the standard Clinical & Laboratory Standards Institute (CLSI) approved broth microdilution (BMD) methods with cation-adjusted Mueller-Hinton Broth (caMHB). Resistance was determined as per CLSI guidelines [27] except for tigecycline and fosfomycin where The European Committee on Antimicrobial Susceptibility Testing (EUCAST) (Version 7.1, 2017) (see http://www.eucast.org) guidelines were implemented. Categorisation of drug resistance level was determined through guidelines previously outlined [28].

### DNA extraction

DNA was extracted from overnight cultures using the DNeasy Blood and Tissue Kit (Qiagen) with the additional enzymatic lysis buffer pre-treatment as per manufacturer’s instructions. DNA was quantified with Qubit^®^3.0 (ThermoFisher Scientific).

### DNA library preparation and sequencing

Library preparation was performed using the Nextera XT kit (Illumina) with 1 ng input of DNA as per manufacturer’s instructions. Quality of libraries were checked using a 2100 Bioanalyzer (Agilent Technologies). Libraries were sequenced on an Illumina MiSeq with 300 bp paired-end sequencing reads and >100X coverage per sample.

### Sequencing analysis

Paired-end reads were trimmed with Trimmomatic [29] and assembled using SPAdes [30]. The Rapid Annotation using Subsystem Technology (RAST) was utilized to annotate assembled genomes [31]. Assemblies were also uploaded to the Centre for Genomic Epidemiology (CGE) to identify sequence types (STs) (MultiLocus Sequence Typing Server 1.8 [32]) and acquired antibiotic resistance genes (ResFinder 2.1 [33]). A neighbor-joining tree was constructed using the 2358 *Klebsiella pneumoniae/quasipneumoniae/variicola* genes known to form the core genome MLST (cgMLST) using Ridom SeqSphere+ v4.0.1 software [34]. The cgMLST was compared against complete assemblies of ST11 (HS11286), ST147 (MS6671), ST258 (NJST258_1, NJST258_2) and a reference for *K. quasipneumoniae*, ATCC 700603 [25, 35–37].

### Variant detection

Alterations both in and flanking the genes *pmrAB*, *phoPQ* and *mgrB* were examined and sequence reads of all strains were aligned to the assembly of 20_GR_12, a polymyxin-susceptible ST258 strain with the least number of contigs, using BWA-MEM [38]. The alignment was analyzed through FreeBayes [39] to identify single nucleotide and small indel variation, using a diploid analysis in order to identify potential heterogeneity. Sites with more than 20% of reads mapping to the minor allele were considered potentially heterogeneous. The effects of variations were determined by snpEff [40]. The impact on protein sequence was further confirmed by the Protein Variation Effect Analyzer (PROVEAN) [41]. For the analysis of large chromosome changes, the gene sequences including 300 bp flanking were extracted from the assemblies. A multiple alignment of each gene was constructed from the pair-wise alignment to the longest gene sequence.

### Insertion sequence element validation

ISFinder [42] was used for the identification of insertion sequence (IS) elements. To confirm disruptive IS elements, *mgrB* was amplified with primers displayed in Table S1 via 2X Phusion HF master mix (Invitrogen) under the following cycling conditions: 98 °C 10 seconds, 50 °C 30 seconds and 72 °C 60 seconds (35X). Amplicon identity was validated via Sanger sequencing.

### Complementation assays

The contribution of variants to resistance was validated through complementation assays as previously described [43]. Briefly, genes (Table S1) were amplified from a polymyxin-susceptible isolate, 20_GR_12, and cloned into the pCR-BluntII-TOPO vector via the Zero Blunt TOPO PCR cloning kit (Invitrogen). Chemically competent *E. coli* TOP10 cells were transformed and selected by the addition of 50 mg/L kanamycin in MHA. Isolation of plasmids were via the PureLink™ Quick Plasmid Miniprep Kit (Invitrogen) and transformed into KP strains via electroporation (25 μF, 200 Ω, 1.25 kV/cm) with a Gene Pulser (Bio-Rad Laboratories). Selection was accomplished through supplementation of ≥500 mg/L zeocin in MHA plates. Transformed colonies (n=≥2) were acquired and placed in MHB containing 1500 mg/L zeocin and 1 mM isopropyl *β-D-1-* thiogalactopyranoside (Sigma Aldrich). If polymyxin susceptibility was not restored upon complementation, genes harboring mutations were further amplified and introduced into 20_GR_12. To discern the impact of additional mutations in *phoPQ* and *pmrB* segregating with disrupted *mgrB*, mutant genes were introduced into a polymyxin-resistant isolate only harboring an IS element *mgrB* disruption, 7_GR_13. Controls included transformation of WT genes into 20_GR_12, sequencing of amplicon prior to introduction in vector and KP transformed strains undergoing a plasmid extraction and further PCR of the multiple cloning site. Antimicrobial testing against polymyxin B were conducted as described above.

## RESULTS

### Characterization of clinical isolates

KP isolates were all characterized in the hospital microbiology facility using VITEK^®^2. Several discrepancies were detected between VITEK^®^2 and broth microdilution (BMD) results (Table 1, Table S2), predominantly the level of resistance towards aminoglycosides, tetracyclines, fosfomycin and tigecycline. A major dissimilarity was polymyxin susceptibility in 6_GR_12 (sensitive in BMD, resistant in VITEK^®^2) and resistant in 23_GR_13 (resistant in BMD, sensitive in VITEK^®^2). Polymyxin resistance was identified in 19 of the isolates. An abundance of acquired resistance genes (Table 2) were detected and this presence corresponded to the antimicrobial testing phenotype. This analysis did not identify *mcr-1* in these strains. Only 18_GR_14 and 19_GR_14 were not identified as extended-spectrum beta-lactamase producers amongst the polymyxin-resistant strains. Consequently, all polymyxin-resistant strains that harbored non-susceptibility to at least one antibiotic in 15 or more of the 17 antimicrobial categories hence were defined as XDR.

**Table 1.**
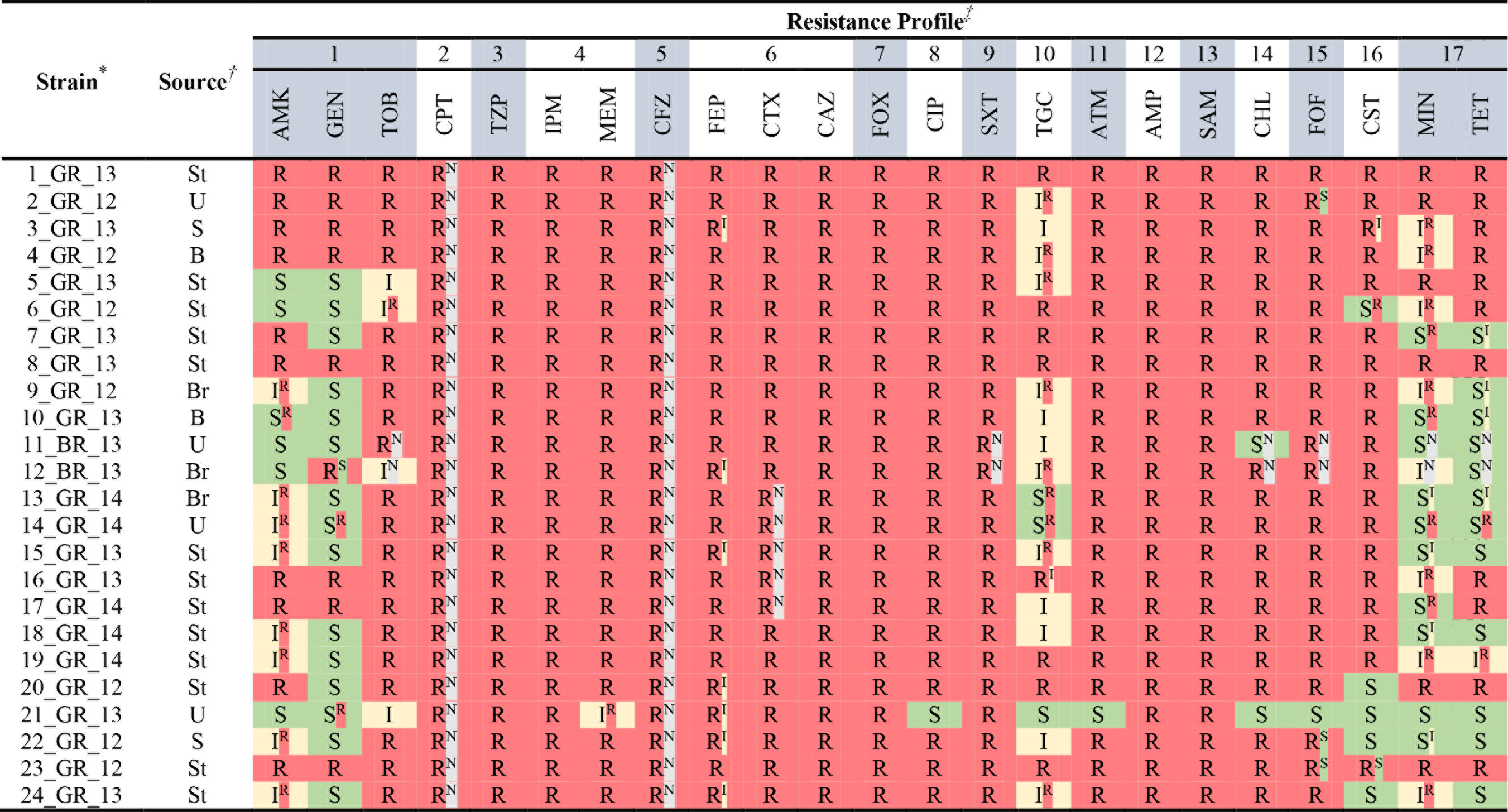
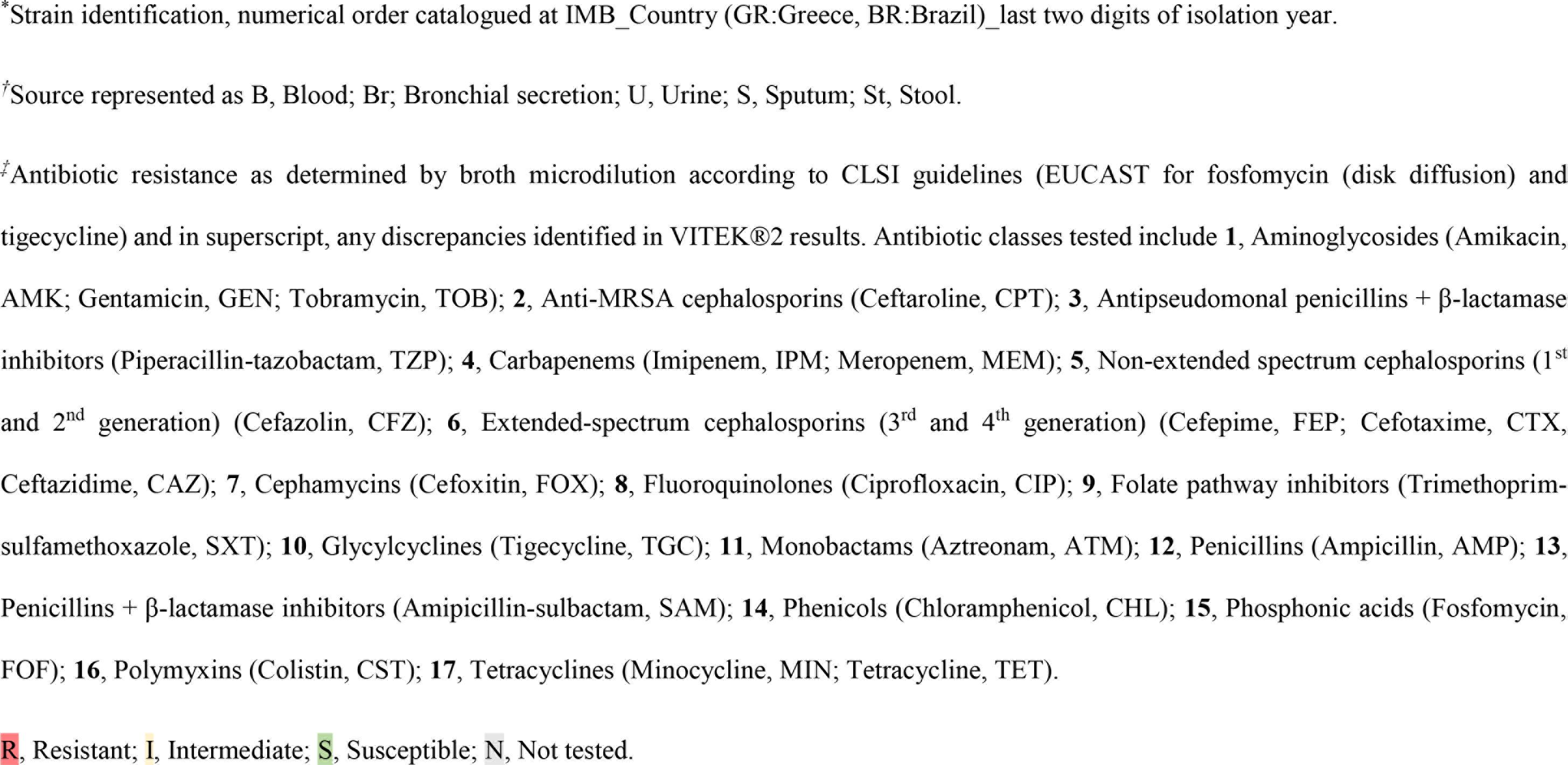
Broth microdilution and VITEK^®^2 antimicrobial testing for the 24 clinical isolates.

**Table 2.**
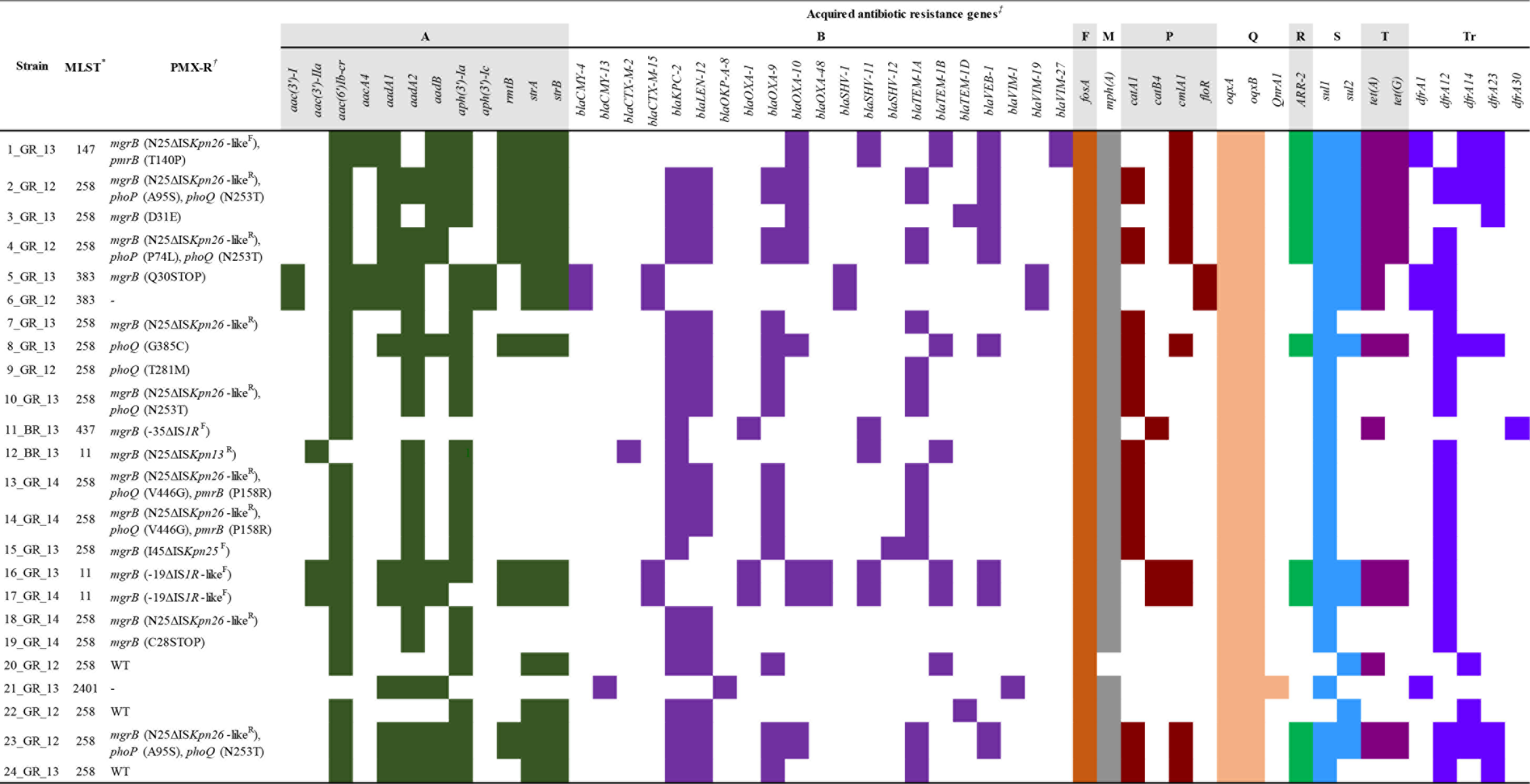
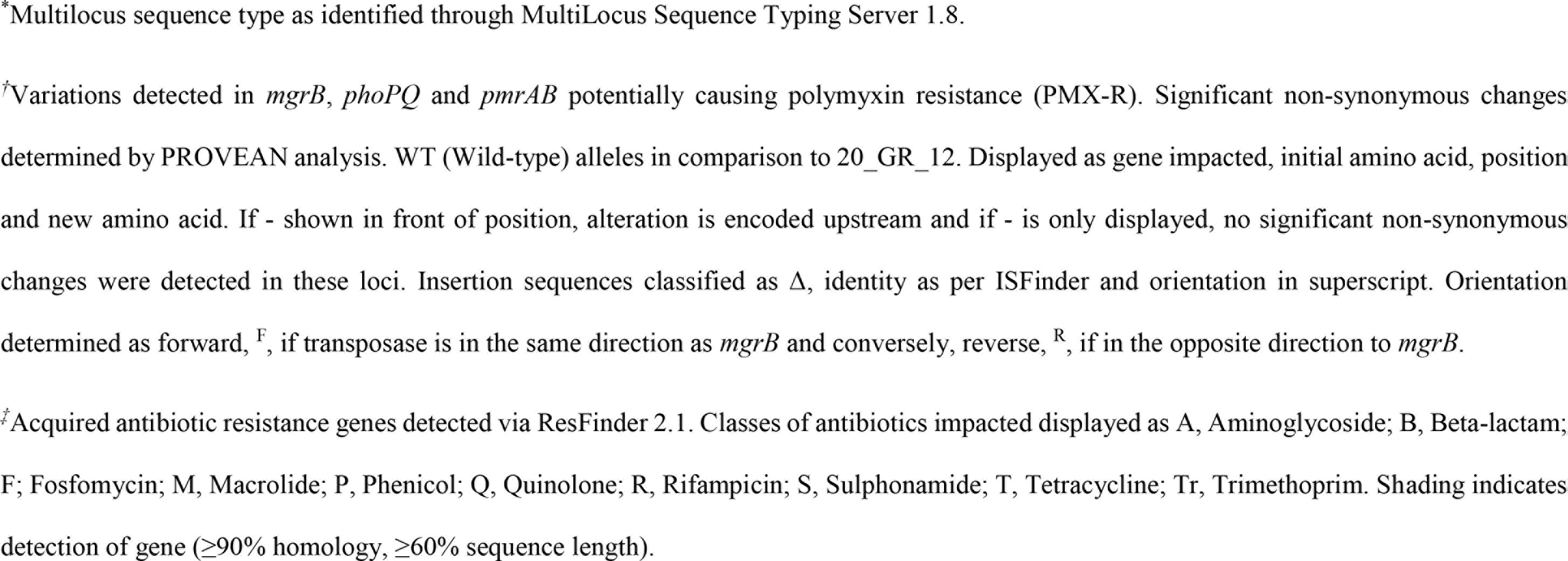
Potential mutations contributing to polymyxin resistance and acquired resistance genes.

### Sequence type determination

Two thirds of the Greece clinical strains were found to belong to ST258 and the remaining were ST11, ST147 or ST383 (Table 1). While 5_GR_13 and 6_GR_12 were both ST383, only 5_GR_13 was resistant to polymyxin. Among the two strains from Brazil, 11_BR_13 was ST437 and 12_BR_13 was ST11. 21_GR_13 had a profile previously undefined and has been newly designated ST2401. Further cgMLST studies were conducted on the isolates using complete assemblies of reference genomes for ST11 (HS11286), ST147 (MS6671) ST258 (NJST258_1, NJST258_2) and KQ (ATCC 700603) (Fig. 1). For the ST258 isolates, these were more similar to NJST258_2 rather than NJST258_1. Within this cluster, 7_GR_13, 9_GR_12 and 24_GR_13 were closely related (≤15 allelic changes). Similarly, grouped together were 2_GR_12 and 23_GR_12; 3_GR_13 and 22_GR_12; 13_GR_14 and 14_GR_14; and 18_GR_14 and 19_GR_14. In ST11, 16_GR_13 and 17_GR_14 harbored only 3 allele differences and the Brazilian isolate, 12_BR_13, had 206 variants apparent. ST383 isolates 5_GR_13 and 6_GR_12 only exhibited 1 allele change. ST147 1_GR_13 was not clonal to the previous pandrug resistant KP, MS6671. Clustering analysis revealed 21_GR_13 as *Klebsiella quasipneumoniae* (KQ) and diverged with reference genome ATCC 700603 (ST489).

**Fig. 1.**
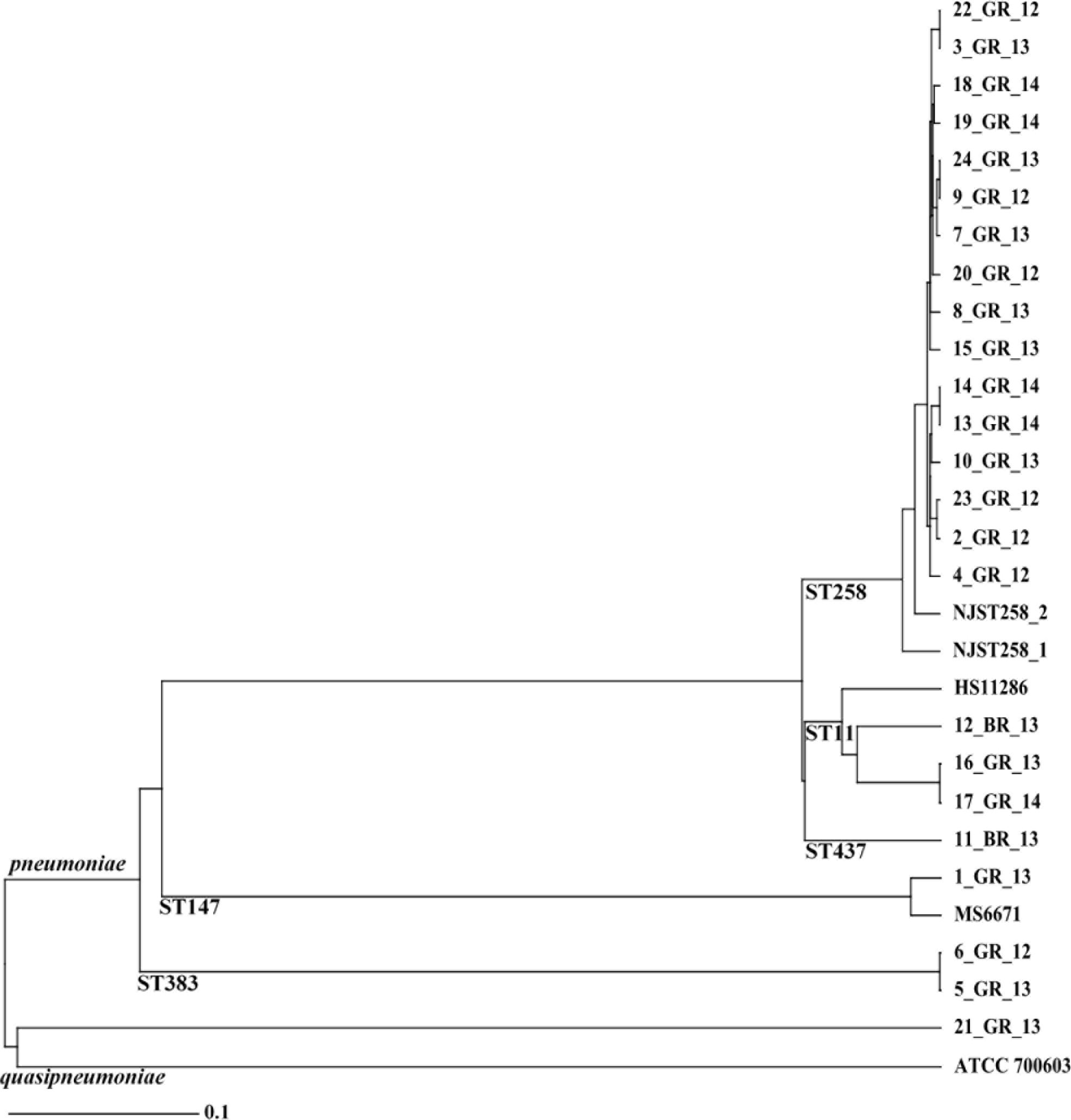
Neighbor-joining tree of core genome MLST of 24 *Klebsiella* clinical isolates. Clustering of sequence type (ST) indicated at base of diverging branch. cgMLST compared to completed assemblies including ATCC 700603 (*K. quasipneumoniae*), HS11286 (ST11), MS6671 (ST147) and NJST258_1 and NJST258_2 (ST258)

### MgrB disruption

Seventeen of the nineteen polymyxin-resistant strains exhibited either missense mutations, nonsense mutations or IS elements in *mgrB* (Table 3). Both 5_GR_13 and 19_GR_14 harbored a truncation while an amino acid change, D31E, was apparent in 3_GR_13. IS element disruption was prevalent in 53% of strains and commonly an IS5-like element was integrated at nucleotide position 75 (Fig. S1). Sanger sequencing revealed this element was closely related to IS*Kpn26*, herein known as IS*Kpn26*-like, except for 12_BR_13 which matched IS*Kpn13*. IS*1R* was detected upstream of *mgrB* in 11_BR_13 and an IS1R-like (A>C, 393 bp; C>T, 396 bp) element in 16_GR_13 and 17_GR_14. Strain 15_GR_13 had a deletion of the *mgrB* locus from nucleotide position 133 onwards. The 127 bp flanking region mapped to ISK*pn25* with the transposase in the same orientation as *mgrB*. All 3 of IS*1* element insertions, but only one of the 8 IS*Kpn26*-like element insertions had the transposase in the same orientation as *mgrB*.

### Single, multiple and heterogeneous mutations

Aberrations in genes commonly identified to confer polymyxin resistance in KP include *mgrB*, *phoPQ* and *pmrAB* (Table 2). Several non-synonymous mutations were identified across the isolates however, not all were predicted to be deleterious (Table S3). ST383 contained several mutations in *pmrAB* although only Q30STOP in polymyxin-resistant 5_GR_13 was predicted to have an impact. Similarly, neutral changes in all four of these genes were detected in polymyxin-susceptible KQ strains ATCC 700603 and 21_GR_13. 8_GR_13 and 9_GR_12 harbored a single detrimental missense mutation in *phoQ*. Alterations in *mgrB* were accompanied by one or more missense mutations in *phoPQ* and/ or *pmrB*. Predicted deleterious variants segregating with disrupted *mgrB* included *pmrB* (T140P, P158R), *phoP* (P74L, A95S) and *phoQ* (N253T, V446G), which were commonly in the ST258 lineage. V446G (*phoQ*) and P158R (*pmrB*) were heterogeneous in 13_GR_14 (65% (V446G), 66% (P158R) mutation allele frequency) and 14_GR_14 (52% (V446G) and 57% (P158R) mutation allele frequency). Assembly revealed 23_GR_12 harbored an IS*Kpn26*-like disrupted *mgrB* alongside the intact version with alterations in *phoP* and *phoQ* in 57% of the sample.

### Role of *mgrB* disruptions and presence of heteroresistance via complementation assays

Complementation of the WT gene elucidated the role of these mutations in resistance (Fig. 2). Introduction of pTOPO-*mgrB* restored susceptibility in all resistant isolates with *mgrB* coding mutations or upstream disruptions, with the exception of two strains heterogeneous for the *mgrB* disruption and a *pmrB* coding mutation (13_GR_14 and 14_GR_14) (Fig. 2a). For these two strains, pTOPO-*mgrB* restored susceptibility in zero of three 13_GR_14 colonies and one of three 14_GR_14 colonies. Transformation of 1 out of 3 colonies for both 13_GR_14 and 14_GR_14 strains with pTOPO-*pmrB* restored susceptibility (Fig. 2d) and *mgrB* amplification of these colonies revealed an intact *mgrB* locus (data not shown). Colonies which were reverted on complementation were further passaged 3 times with no antibiotic pressure in order to remove the plasmid and discern if these mutations were contributing to resistance. After passaging, pTOPO-*mgrB* isolates harbored an MIC of ≥64 mg/L whilst pTOPO-*pmrB* colonies were 16 mg/L to confirm two resistant populations in these samples. 23_GR_12 was also observed to have a heterogeneous *mgrB* disruption but did not carry a corresponding *pmrB* mutation however, harbored similar mutations to 2_GR_12 in *phoPQ*. Amplification of *mgrB* identified two of three 23_GR_12 transformed colonies contained the IS element disruption and were reverted to susceptible upon complementation with pTOPO-*mgrB*.

**Fig. 2.**
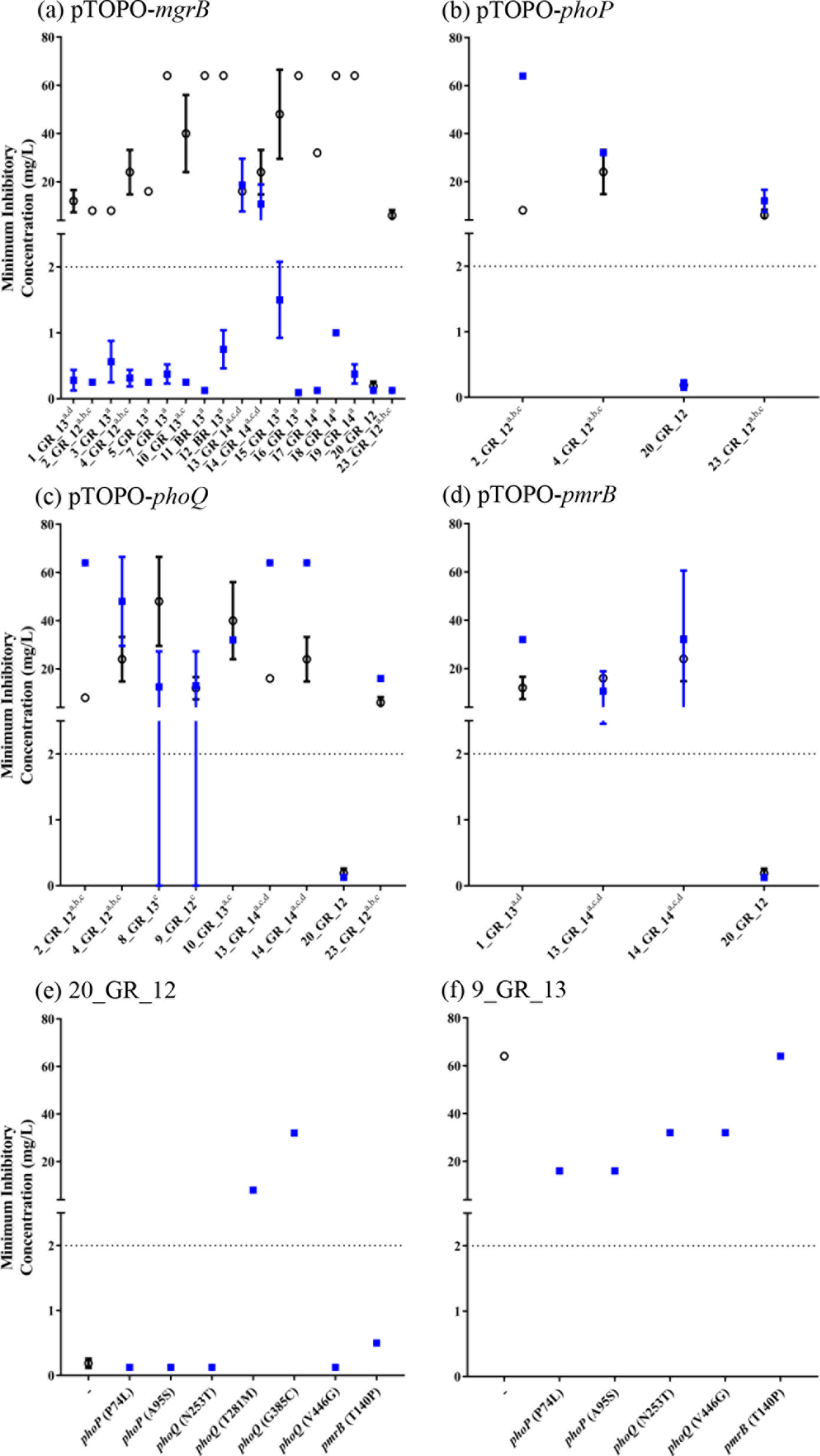
Complementation assays and influence of gene on polymyxin resistance. Polymyxin B MIC measured before (○) and after (◾) complementation of wild-type gene (a) *pTOPO-*mgrB**, (b) pTOPO-phoP, (c) pTOPO-phoQ, or (d) pTOPO-*pmrB* in indicated resistant isolates. (e) Mutated genes complemented into 20_GR_12 (polymyxin-susceptible isolate) to determine if variant induces polymyxin resistance. (f) Complementation of 9_GR_13 (IS element disrupted *mgrB* control) to detect potential suppressor mutations. Strains shown on x axis for (a-d) and superscript indicates variants in genes including *mgrB* (a), *phoP* (b), *phoQ* (c) and *pmrB* (d) that differ from 20_GR_12. For (e, f), the x axis shows the gene complemented with amino acid variation in brackets. Dotted line at 2 mg/L represents the breakpoint for polymyxin B. Values indicate mean±standard deviation where no error bars display no fluctuation in MIC (n≥2 colonies).

### Validation of resistance conferring mutations in *phoQ*

Strains 8_GR_13 and 9_GR_12 harbored a single mutation in *phoQ* potentially conferring resistance (Table 2). When these isolates were transformed with pTOPO-*phoQ*, results remained variable where a lack of growth was present in a susceptible range (MIC: ≤2 mg/L) however, several wells containing high polymyxin B concentrations exhibited growth (Fig. 2c). To resolve this, the mutated gene was introduced into a polymyxin-susceptible isolate, 20_GR_12, and resistance was apparent (Fig. 2e).

### Potential suppressor mutations in *phoPQ*

Several mutations co-segregating with disrupted *mgrB* were detected including *phoP* (P74L, A95S), *phoQ* (N253T, V446G) and *pmrB* (T140P). Complementation of WT genes in these isolates commonly facilitated a ≥2-fold increase in MIC with the exception of 10_GR_13, which had an additional predicted neutral mutation in *phoQ* (A225T) (Table S3, Fig. 2b-d). To evaluate the potential influence of these mutations on polymyxin resistance, mutated genes were placed into a strain only containing the *mgrB* IS element disruption, 7_GR_13 (Fig. 2f). Complementation of the mutant *phoQ* (N253T) decreased the MIC by 2-fold, potentially indicating a partial suppressor mutation. Initially, the *phoQ* (V446G) mutation was anticipated to segregate with the *mgrB* disrupted population in 13_GR_14 and 14_GR_14 however, when *phoQ* was amplified from a colony reverted to susceptible via pTOPO-*mgrB* complementation, the WT*phoQ* was observed (Fig. S3). The *phoQ* (V446G) mutation was successfully amplified from a 14_GR_14 colony containing the *pmrB* (T158R) mutation and upon complementation in 7_GR_13, resulted in a 2fold reduction in MIC. Although this mutation did not segregate with disrupted *mgrB*, it may act as a partial suppressor mutation when a resistance conferring mutation is present in *pmrB*. Mutations in *phoP* (P74L, A95S) reduced the MIC in 7_GR_13 by >4-fold which identifies these as partial suppressor mutations. Complementation of mutant *pmrB* (T140P) into 7_GR_13 did not lead to an observable corresponding reduction in MIC however, once transformed into 20_GR_12, a 2-fold increase in MIC was apparent (Fig. 2e).

## DISCUSSION

Polymyxin resistance in XDR-KP is of grave concern given that this is a last-line antibiotic, and is increasingly prevalent in countries such as Greece and Brazil [10, 12–14, 44]. We evaluated the genetic basis of polymyxin resistance in a series of Greek and Brazilian clinical isolates from patients in 2012 to 2014 and found alterations in genes *mgrB*, *phoPQ* and *pmrAB*.

Inactivation of *mgrB* was highly prevalent in these strains with an IS*Kpn26*-like element being the predominant cause of resistance, as confirmed by complementation restoring susceptibility in all isolates. Several other studies have observed an IS5-like element integration in the same position, including reports from Greece, Italy, France, Turkey and Colombia [8, 9, 45, 46]. The *ISKpn26*-like element resembled the same sequence from Greece isolates previously described [46]. We identified that this mutation still persisted in 2014, after being first detected in 2012 [9]. Disruptions in *mgrB* including the IS*Kpn26*-like forward insertion at nucleotide 75 in ST147, IS*Kpn13* integration at nucleotide 75 in ST11 and ISK*pn25* in the ST258 lineage have yet to be reported. We identified *IS1R* or IS1R-like elements positioned upstream of *mgrB* in 3 isolates (11_BR_13, 16_GR_13, 17_GR_14) which were reverted upon complementation indicating an impact on the promoter region.

Truncations identified at position 28 and 30 of *mgrB* have been previously detected, although these were identified in differing STs indicating mutations potentially have arisen independently in Greece [21, 47]. Complementation restored susceptibility to polymyxins for these mutations and this study further revealed the amino acid change D31E in 3_GR_13 to be a resistance conferring alteration. These findings support the notion that intact MgrB is required to confer negative feedback on PhoPQ [8]. The inactivation of *mgrB* is prevalent in polymyxin-resistant KP and may arise owing to its capacity to promote virulence and further attenuate the early host defence response, with little or no fitness cost [48].

Single predicted detrimental mutations were observed in the *phoQ* histidine kinase region, critical for phosphorylation and interaction with *phoP*, in 8_GR_13 (G385C) and 9_GR_12 (T281M). The G385C mutation had previously been reported, [21] however in a differing ST. Complementation revealed an inconsistent MIC for these strains, although when a polymyxin-susceptible isolate was transformed with the mutated gene, full resistance was restored. Dominance of mutated *phoQ* has recently been highlighted and these results may imply the inability of pTOPO-*phoQ* to override the resistance caused by these mutations [49].

Several non-synonymous changes were identified to be not deleterious according to PROVEAN analysis. Notably, these were abundant in KQ strains ATCC 700603 [37] and 21_GR_13. This was further identified in KP ST383 lineages and PROVEAN detected these neutral changes. These mutations represent lineage specific alterations, however, this does not negate the possibility of previously resistance conferring alterations being acquired in these loci with subsequent reversion mutations to give rise to a susceptible phenotype.

Heterogeneity was apparent in several isolates. In near equal ratios, 13_GR_14 and 14_GR_14 possessed the IS*Kpn26*-like *mgrB* disruption and a new alteration conferring resistance in *pmrB*, P158R as determined by complementation. 23_GR_12 consisted of approximately half the reads mapping to the undisrupted genes and the other to the IS*Kpn26*-like strain with several additional predicted deleterious mutations. This heterogeneity may explain the initial clinical detection for this isolate to be polymyxin-susceptible.

Several isolates harboring IS*Kpn26*-like element disrupted *mgrB* were accompanied by mutations in *phoPQ* and/ or *pmrB*. These changes were present in ≥98% of reads to render the involvement of heterogeneity unlikely. Once complemented, an increase in resistance was commonly recorded. This potentially reflects partial suppressor mutations as strains which solely possessed this IS element disruption commonly exhibited a heightened MIC of ≥64 mg/L. One variant segregating with this disruption included *pmrB* T140P. This had formerly been identified in an ST258 lineage but even when the resistant gene was complemented, the MIC increased by 2-fold but was not defined as clinically resistant [21, 50].

When mutated *phoP* or *phoQ* were introduced into the *mgrB* disrupted isolate, a reduction in MIC was apparent. The involvement of additional mutations in PhoPQ to influence the level of polymyxin resistance has yet to be reported in KP. Previous research by Miller *et al* [51] determined additional mutations in PhoPQ altered polymyxin resistance in *Pseudomonas aeruginosa*. This prior study describes *phoP* mutations with the capacity to partially or fully suppress resistance-causing mutations in *phoQ*. These mutations in *phoP* were near or within the DNA binding site which differs to our results, where the alterations are impacting the response regulatory region that interacts with PhoQ. Conversely, all mutations partially suppressing the MIC were identified to be targeting the HAMP and histadine kinase component of PhoQ. These were in regions similar to revertant *P. aeruginosa* strains identified by Lee and Ko [52]. We postulate these mutations are perturbing the critical transfer of phosphoryl groups from the histadine kinase of PhoQ to PhoP and subsequent *pmrD* expression. Whether these mutations constitute a fitness advantage due to the reduction of metabolism required for the production of LPS modifications is yet to be discerned. Furthermore, due to variability in some of the complementation data, a knockout *phoPQ* background and introduction of genes that are potential suppressor mutations is required.

Rapid and accurate detection of mutations attributed to polymyxin resistance remains a longstanding burden. Our research has contributed to the current understanding of the dissemination and evolution of this resistance in KP. Although the sample size is limited, this study highlights several issues arising from solely interrogating genomes for resistance detection including ST specific non-synonymous changes, and heterogeneity. The study provides the first potential report of suppressor mutations for polymyxin resistance. Through complementation assays, we have discerned the role of these modifications and have identified resistance-causing alterations that can be monitored during future genome-based diagnostics.

## Acknowledgements

We thank Dr Aurélie Jayol and Professor Patrice Nordmann for providing their complementation assay methodology. The authors thank Maite Amado and Angela M. Kavanagh for technical support with susceptibility assays and Rhia Stone for the quality control of antibiotics. We acknowledge the sequencing services provided by the Australian Genome Research Facility. We thank the team of the curators of the Institut Pasteur MLST system (Paris, France) for importing novel alleles, profiles and/or isolates at http://bigsdb.web.pasteur.fr.

## ABBREVIATIONS

BMD, Broth microdilution; caMHB, cation-adjusted Mueller-Hinton broth; CLSI, Clinical & Laboratory Standards Institute; EUCAST, The European Committee on Antimicrobial Susceptibility Testing; IS, Insertion sequence; KP, *Klebsiella pneumoniae*; KQ, *Klebsiella quasipneumoniae*; MHA, Mueller-Hinton agar; MHB, Mueller-Hinton broth; MIC, Minimum Inhibitory Concentration; MLST, Multi-locus sequence type; pEtN, phosphoethanolamine; PROVEAN, Protein Variation Effect Analyzer; ST, Sequence type; XDR, Extensively drug resistant; XDR-KP, Extensively drug-resistant *Klebsiella pneumoniae*.

